# Likelihood-based evaluation of character recoding schemes for phylogenetic analysis

**DOI:** 10.1101/2025.10.15.682473

**Authors:** Tae-Kun Seo, Jeffrey L. Thorne

**Author notes:** Correspondence (T-K S).

## Abstract

Character recoding is a common practice in evolutionary studies. For example, phylogenies can be inferred from protein-coding DNA sequences via 61-state codon substitution models. However, often the inference is instead done by adopting 20-state amino acid replacement models that do not explicitly consider synonymous substitution. When there is substantial heterogeneity of amino acid frequencies among sites and/or among lineages, another sort of character recoding is sometimes performed. In these cases, one option is to reduce the state space of the model by placing each of the 20 amino acids into one of a relatively small number (e.g., 6) of groups of amino acids and then modeling only how group membership changes at a site over evolutionary time. Unfortunately, these kinds of character recoding schemes are prone to reducing the amount of available evolutionary information. Here, we provide a likelihood framework to statistically assess recoding schemes. Although we concentrate on the recoding of 61-state codon substitution models into 20-state amino acid replacement models, the general approach is also relevant to other recoding schemes such as those that recode 20-state models into 6-state models.

## INTRODUCTION

Recoding character states is frequently conducted in phylogenetic analysis. When saturation of synonymous substitutions is suspected, 61-state codons are sometimes recoded (i.e., translated) into 20-state amino acids and phylogenetic analysis is then performed on the basis of amino acid sequences. Similarly, when compositional heterogeneity of amino acids among sites and/or among lineages is suspected, 20-state amino acid characters can be recoded into fewer states (e.g., 6 states) with the hope that compositional heterogeneity will be mitigated in the smaller dimensional space (e.g., see Hernandez and Ryan 2021). When saturation of nucleotide transition substitutions is a concern, the four nucleotide types can be recoded as being either purine or pyrimidine and then a two-state evolutionary model that reflects only transversion substitutions can be applied (Phillips and Penny 2003). Despite their diverse natures, these recoding schemes all ignore within-group substitutions that do not result in state changes in the recoded character space. Here, we concentrate on the within-group substitutions that are ignored when recoding is performed.

A typical recoding scheme of amino acids is a 6-state recoding that was derived from the Dayhoff model of amino acid replacement (Dayhoff et al. 1978) and many variations have been suggested (Kosiol et al. 2004; Susko and Roger 2007). Although 6-state amino acid recoding has often been adopted for phylogenetic analysis (e.g., see relevant references listed in Hernandez and Ryan 2021), its effectiveness is questionable because some research has suggested that unrecoded sequence data performs better than recoded data (Hernandez and Ryan 2021). An alternative opinion is that recoding sequences to a lower dimension of character states can either increase or decrease phylogenetic accuracy (Foster et al. 2023).

A continuous-time discrete-state Markov process on the original state space is transformed by recoding to a stochastic process that operates in a smaller (recoded) state space. Lumpability (Kemeny and Snell 1976) refers to the situation where the process on the recoded character space is also Markovian. When the lumpability property is satisfied, the probability of ending at a specific recoded character state can depend on the specific recoded state that was occupied at the beginning of the process but cannot additionally depend on which state of the original (bigger) state space was occupied at the beginning.

Some character state recodings have the lumpability property whereas others do not. Lumpability tests for DNA characters have been developed for the phylogenetic analysis of two sequences (Vera-Ruiz et al. 2014) and multiple sequences (Vera-Ruiz et al. 2022). When character recoding violates lumpability, a Markov model on the recoded states will be difficult to reconcile with a Markov process on the original states. However, the recoded Markov model can still be viewed as an approximation of change among the recoded character states.

Recoding character states should be a thoughtful decision. Smaller numbers of character states can yield computational advantages, but they can also lead to loss of evolutionary information. Even for highly divergent sequence data, simply adopting a 20-state amino acid model should not be done without carefully investigating how much evolutionary information is being lost by ignoring synonymous substitutions (Seo and Kishino 2008). Likewise, recoding nucleotides into purines and pyrimidines and then using a 2-state “RY” model rather than a 4-state nucleotide substitution model runs the risk of ignoring evolutionary information from transition substitutions. In addition, choosing recoding due to concerns about compositional heterogeneity among sequences may be ill-advised without statistically assessing whether the potential drawbacks of lost evolutionary information are warranted.

Determining whether to adopt a recoding scheme is a problem of model selection. Statistical approaches for comparing models in phylogenetics generally derive a conclusion such as “Model A is better in Situation A and Model B is better in Situation B.” For example, it is widely accepted that amino acid models are better when protein-coding sequences are highly divergent, whereas DNA or codon models are better when protein-coding sequences have less divergence (e.g., see Seo and Kishino 2008, 2009; Miyazawa 2013; Simmons 2017). Unfortunately, it can be unclear whether a data set of interest actually corresponds to “Situation A” or “Situation B” or something intermediate between these situations. Conclusions that are derived from simulation studies are necessarily limited to the scenarios that were explored. While exceptions exist (e.g., Venkat et al. 2018; Abadi et al. 2019), the focus of most simulation studies pertaining to recoding has been on accuracy of tree topology estimation and not on inference of other characteristics of potential interest (e.g., strength of natural selection, branch lengths, evolutionary rates, ancestral states).

Here, we suggest a likelihood framework to evaluate the effect of recoding schemes. Specifically, we attempt to measure the difference in information between the original and recoded character spaces. Under this likelihood framework, substitution models for recoded characters and unrecoded characters are comparable via conventional model comparison criteria such as AIC (Akaike Information Criterion; Akaike 1974), BIC (Bayesian Information Criterion; Schwarz 1978), or a likelihood ratio test (LRT). Our approach is a generalization of the Seo and Kishino (2008) approach for comparing amino acid replacement and codon substitution models. We emphasize that quantitative investigation should be made for selecting recoding schemes.

## THEORY

### Assumptions and notation

As noted in the INTRODUCTION, a common feature of all pairs of original and recoded character spaces is that within-group substitutions are ignored in the recoded character space. Although our approach is applicable to other recoding schemes, we describe it using the case of 61-state codon substitution models recoded as 20-state amino acid replacement models.

Denote *a*_*i*_ as an amino acid type that is indexed with the integer *i* (1 ⩽*i*⩽ 20) and *a*(*r*) as the amino acid type encoded by codon *r* (1⩽ *r*⩽ 61). For a given *a*(*r*), there is an index *i* such that *a*_*i*_ *= a*(*r*). Denote *π*_*r*_ and *π*_*a*(*r*)_ as the stationary frequencies of codon *r* and amino acid *a*(*r*), respectively. Under the framework of Maximum Likelihood (ML) estimation, stationary frequencies should be estimated by maximizing the likelihood function. However, because ML estimation of codon frequencies can be computationally intensive, codon frequencies are typically estimated from empirical frequencies of the data being analyzed. Here, we follow this convention and infer stationary frequencies of amino acid types by summing the stationary frequencies of all codons that encode the corresponding amino acids,

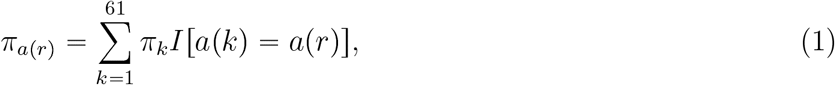

where *I*[·] is defined as 1 if the condition within the brackets is satisfied and 0 otherwise.

### Overview of previous codon substitution models

The GY94 codon model (Goldman and Yang 1994) defines instantaneous transition rates from codon *r* to *s* with *r* ≠ *s* as

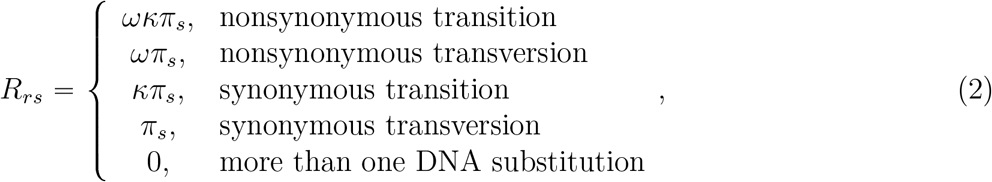

where *κ* differentiates transition and transversion rates and *ω* differentiates nonsynonymous and synonymous rates. The diagonal elements of the rate matrix are the negative row sums: {*R*_*rr*_ = ∑_*s*:*r≠s*_ *R*_*rs*_}. This definition of diagonal elements also applies to the diagonal elements of 20-state and 4-state models that will be discussed below.

Conventional 20-state amino acid models (e.g., Dayhoff et al. 1978; Jones et al. 1992; Adachi and Hasegawa 1992; Whelan and Goldman 2001; Le and Gascuel 2008) define the instantaneous rate from amino acid *a*_*i*_ to amino acid *a*_*j*_ with *a*_*i*_ ≠*a*_*j*_ as

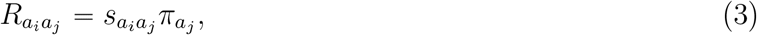

where the 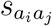’s are exchangeability coefficients (ECs) derived from empirical observation of amino acid changes among a large number of sequences. For the convenience of calculation, we normalize ECs via

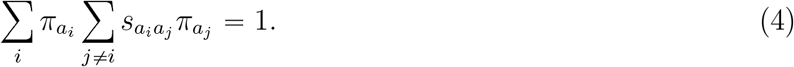

Using the 20-state general time-reversible model of (3), Seo and Kishino (2008) developed the SK-P1 61-state codon model which has rates from codon *r* to *s* (*r* ≠*s*) as

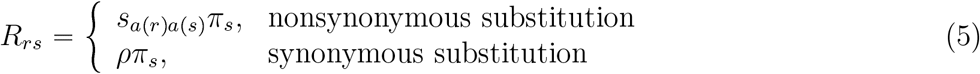

where the *s*_*a*(*r*)*a*(*s*)_ terms are from (3) and the parameter *ρ* represents the relative occurrence of synonymous codon substitutions. Whereas multiple nucleotide changes are not allowed during infinitesimal time in the GY94 model, they are allowed in the SK-P1 model. For simplicity of description, we do not normalize the SK-P1 rate matrix. The SK-P1 transition probabilities for time *t* are

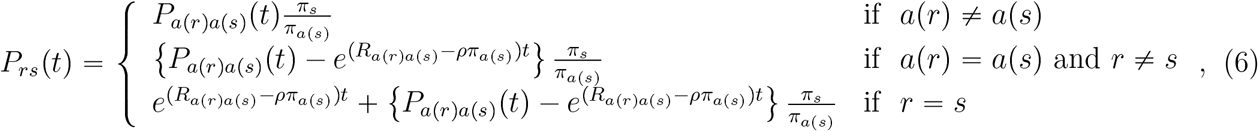

where *P*_*a*(*r*)*a*(*s*)_(*t*) is a transition probability of the conventional 20-state model of (3) for time *t*.

Seo and Kishino (2008) showed that the conventional 20-state model of (3) and the 61-state SK-P1 model become mathematically equivalent when *ρ =* ∞ in (5). That is, ‘ignoring synonymous substitution’ is equivalent to ‘assuming synonymous saturation’ (i.e., *ρ=* ∞ with SK-P1). The special case of the SK-P1 model with *ρ =* ∞ is referred to as the SK-P0 model. Under the SK-P0 model, the transition probability from codon *r* to codon *s* is

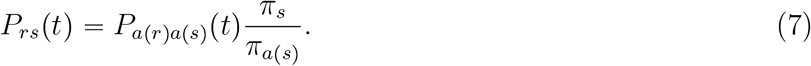

This relationship leads to the following equality between the likelihoods of the 61-state SK-P0 model and 20-state amino acid models,

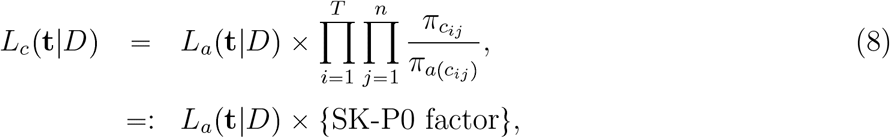

where the ‘=:’ notation implies the right-hand side equation is defined as the left-hand side equation and where *T, n*, and *c*_*ij*_ respectively represent the number of taxa, the length in codons, and the codon at the *j*th site of the *i*th taxon. The *L*_*c*_(·) and *L*_*a*_(·) terms are respectively likelihoods under the 61-state SK-P0 model of (5) with *ρ =* ∞ and under the 20-state amino acid model of (3). The second term of the right-hand side of (8) will be referred to as the ‘SK-P0 factor’ and was also referred to as an ‘adaptor function’ during projection of an amino acid model to a codon model (Whelan et al. 2015).

A widely-employed alternative to models that assume all protein positions evolve according to the same amino acid replacement process is a variation that adopts a mixture of amino acid replacement processes across positions (Le et al. 2008). This variation specifies the number of mixture components (i.e., the number of possible different amino acid replacement processes) and the prior probabilities of sites evolving according to each of the mixture components. If one makes the assumption that mixture components do not vary in the proportions per amino acid of each codon type (i.e., the assumption that 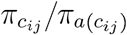 are invariant among mixture components), then the property of (8) still holds. Specifically, the assumption enables the SK-P0 model to be easily modified to mixtures of codon substitution processes with the likelihood of this modified mixture model being equal to the SK-P0 factor multiplied by the likelihood of the amino acid replacement mixture model.

Although the SK-P1 model is an appealing bridge between 20-state amino acid replacement models and 61-state codon substitution models, it tends to provide a poorer model fit than widely-used codon models (Seo and Kishino 2008). One of the reasons for the poor performance of the SK-P1 model is that it ignores important features such as the number of codon positions that change in a codon substitution and whether codon changes are transitions or transversions. In the next subsection, we address this shortcoming with new models.

The SK-P2 codon model (Seo and Kishino 2008) modifies the GY94 model by replacing the *ω* parameter with *s*_*a*(*r*)*a*(*s*)_’s from the amino acid model of (3). For *r* ≠*s*, the SK-P2 model has rates

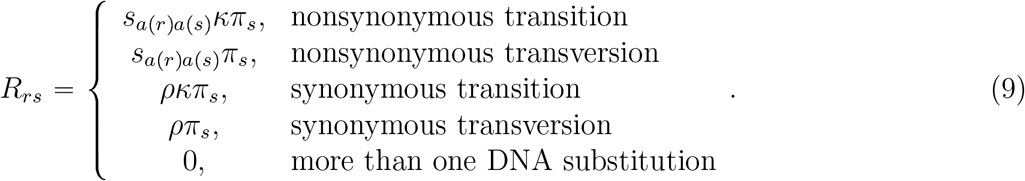

As in the SK-P1 model, the *ρ* parameter is assigned to synonymous substitution. The GY94 model can be considered the special case of the SK-P2 model with all *s*_*a*(*r*)*a*(*s*)_ terms set to 1 and *ρ =* 1/*ω*.

A weighted average of the *s*_*a*(*r*)*a*(*s*)_ values of the SK-P2 model can be obtained as

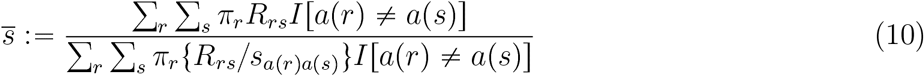

and the ratio 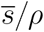 can be used to measure the selection strength. The interpretation of 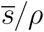 is compatible with the conventional *ω* and *d*_*n*_/*d*_*s*_ measures of selection strength.

### New codon model: Extending amino acid recoding

Generalizing the idea of the SK-P1 model, we define a new codon model whose instantaneous rates are

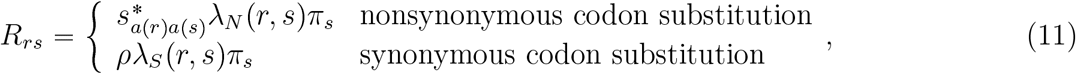

with *λ*_*i*_(*r, s*) for *i* ∈ {*N, S*} being

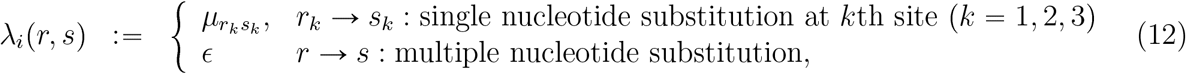

and where 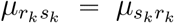 is an EC of the 4-state GTR model (Tavaré 1986) that has *r*_*k*_, *s*_*k*_ ∈ {*A, C, G, T* } and has rates

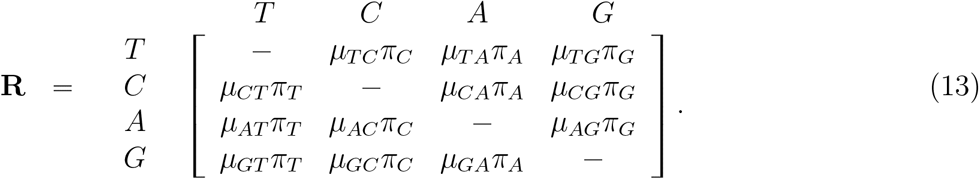

We will consider special cases of the above model that disallow instantaneous multiple nucleotide changes by setting *ϵ =*0, but we will also consider the more general and realistic situation in which *ϵ* ⩾ 0.

One way to specify the 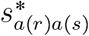 values of (11) is to set them equal to the *s*_*a*(*r*)*a*(*s*)_ from an empirically-derived 20-state amino acid replacement model of (3) and (4). Unfortunately, this approach does not guarantee that the amount of nonsynonymous substitution is equal to the amount of amino acid replacement in the 20-state amino acid model. Another way to set the 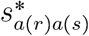 values is to synchronize the nonsynonymous codon substitution rates and the amino acid replacement rates as follows:

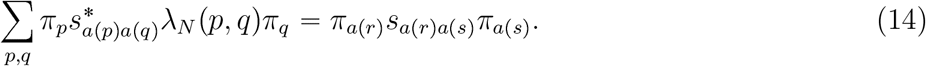

where 1 *⩽ p, q⩽* 61 with *a*(*p*)= *a*(*r*) and *a*(*q*)=*a*(*s*). With this synchronization,

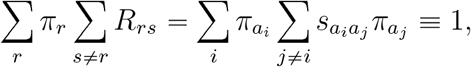

where the second equality comes from normalization of the 20-state amino acid model in (4).

We will refer to the *λ*_*i*_(*r, s*) function by using the name of a corresponding 4-state nucleotide sub-stitution model. Because the 4-state GTR model is employed in (13), we denote *λ*_*i*_(*r, s*) ≡GTR ⊕*ϵ*. When *ϵ =* 0 is explicitly assumed, we denote the function without the ‘⊕*ϵ*’ tag: *λ*_*i*_(*r, s*) ≡GTR. When the HKY model is employed instead of the GTR model (e.g., see RESULTS and DISCUSSION), we denote *λ*_*i*_(*r, s*) HKY⊕ *ϵ*. We note that synchronization of (14) is not possible when *λ*_*N*_ (*r, s*) GTR or HKY, because some amino acid replacements involve only multiple nucleotide substitution. When synchronization of (14) is not possible for *λ*_*N*_ (*r, s*), we should use *s*_*a*(*r*)*a*(*s*)_ in (11)without synchronization.

Various schemes can be employed to define *λ*_*N*_ (*r, s*) and *λ*_*S*_(*r, s*). For example, *λ*_*N*_ (*r, s*) can be constrained to equal *λ*_*S*_(*r, s*). Also, *λ*_*N*_ (*r, s*) and *λ*_*S*_(*r, s*) need not be defined with identical 4-state DNA substitution models. Conventional codon models can be considered special cases of (11). For example, by adopting *λ*_*N*_ (*r, s*) =*λ*_*S*_(*r, s*) ≡ HKY and 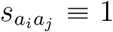, the GY94 model is derived from (11). The special case that is the SK-P1 model has *λ*_*N*_ (*r, s*) =*λ*_*S*_(*r, s*) ≡ 1 where all 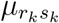and *ϵ* are set to one. Relationships with other special cases are shown in Table 1.

**Table 1:**
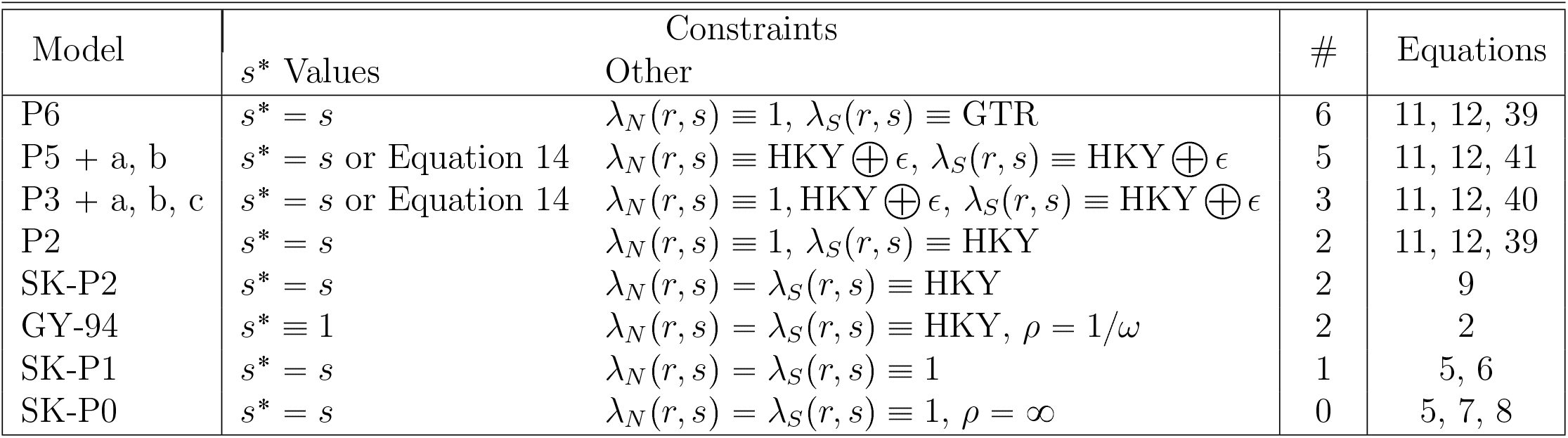
The P6 and P5 models that were defined in RESULTS and DISCUSSION and their special cases. Each row lists a model that emerges from specific constraints on the P6 and P5 parameters. In the 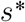 values column, “*s*^*^ *s*” indicates that 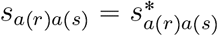 for all combinations of *a*(*r*) ≠ *a*(*s*) whereas entries that are “Equation 14” indicate that the *s*^*^ values are set with Equation 14. The “other” column indicates other constraints on P6 and P5 parameters. The “#” column shows the number of remaining free parameters after constraints are imposed.

| Model | Constraints |  | # | Equations |
|---|---|---|---|---|
|  | s<sup>*</sup> Values | Other |  |  |
| P6 | s<sup>*</sup> = s | λ<sub>N</sub>(r, s) ≡ 1, λ<sub>S</sub>(r, s) ≡ GTR | 6 | 11, 12, 39 |
| P5 + a, b | s<sup>*</sup> = s or Equation 14 | λ<sub>N</sub>(r, s) ≡ HKY ⊕ ϵ, λ<sub>S</sub>(r, s) ≡ HKY ⊕ ϵ | 5 | 11, 12, 41 |
| P3 + a, b, c | s<sup>*</sup> = s or Equation 14 | λ<sub>N</sub>(r, s) ≡ 1, HKY ⊕ ϵ, λ<sub>S</sub>(r, s) ≡ HKY ⊕ ϵ | 3 | 11, 12, 40 |
| P2 | s<sup>*</sup> = s | λ<sub>N</sub>(r, s) ≡ 1, λ<sub>S</sub>(r, s) ≡ HKY | 2 | 11, 12, 39 |
| SK-P2 | s<sup>*</sup> = s | λ<sub>N</sub>(r, s) = λ<sub>S</sub>(r, s) ≡ HKY | 2 | 9 |
| GY-94 | s<sup>*</sup> ≡ 1 | λ<sub>N</sub>(r, s) = λ<sub>S</sub>(r, s) ≡ HKY, ρ = 1/ω | 2 | 2 |
| SK-P1 | s<sup>*</sup> = s | λ<sub>N</sub>(r, s) = λ<sub>S</sub>(r, s) ≡ 1 | 1 | 5, 6 |
| SK-P0 | s<sup>*</sup> = s | λ<sub>N</sub>(r, s) = λ<sub>S</sub>(r, s) ≡ 1, ρ = ∞ | 0 | 5, 7, 8 |

### Rate matrix structure of new model

Here, we focus on setting *λ*_*N*_ (*r, s*) 1 in (11) because this setting yields the appealing bridge between 61-state and 20-state models that the SK-P1 model has while also permitting a less constrained treatment of synonymous substitutions than found in SK-P1. In addition to setting *λ*_*N*_ (*r, s*) ≡ 1 in (11), consider *λ*_*S*_(*r, s*) ≡GTR so that *ϵ* is explicitly set to zero. Because *λ*_*N*_ (*r, s*) ≡1, the setting yields 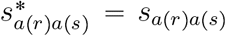 according to (14). Denote the {61 × 61} rate matrix of this model as **R**^(*NS*)^. This rate matrix can be separated into nonsynonymous and synonymous parts to yield two rate matrices,

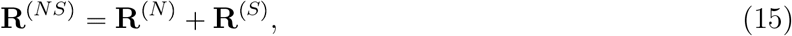

where the nonzero entries in **R**^(*N*)^ (**R**^(*S*)^) correspond to the nonsynonymous (synonymous) changes of **R**^(*NS*)^.

For the convenience of notation, we arrange the order of codons by grouping codons that encode the same amino acid and by ordering these codon groups to match the amino acid ordering in a 20-state amino acid model (see Seo and Kishino 2008). For example, if *a*_1_ of (3) is arginine (Arg), we place the 6 arginine codons into the first part of the **R**^(*NS*)^ matrix. The function *λ*_*S*_(*r, s*) of (12) defines a {6 × 6} synonymous transition rate matrix among these 6 codons. When *λ*_*S*_(*r, s*) ≡ GTR,

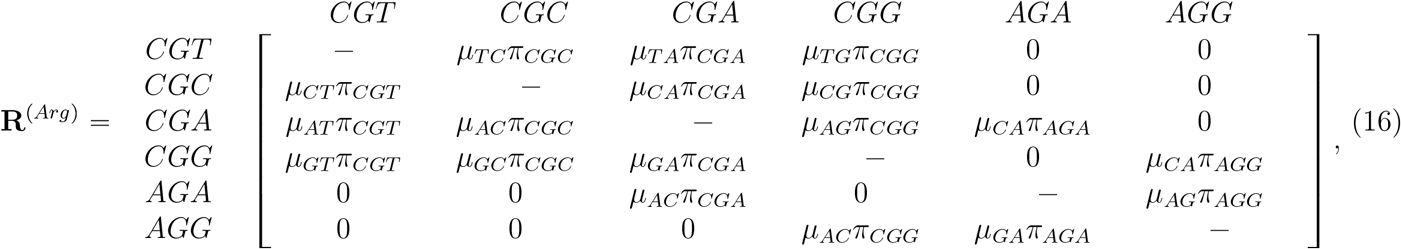

where the diagonal entries are the negative row sums of the five off-diagonal entries in each row.

Then, **R**^(*S*)^ of (15) can be represented as a diagonal block matrix,

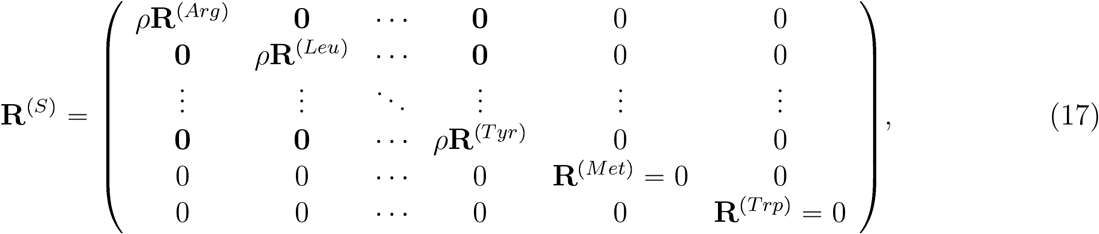

where the (arbitrary) order of encoded amino acids along the diagonal is arginine, leucine, and so on. Because the last two amino acids in (17) (i.e., methionine and tryptophan) are each encoded by a single codon, synonymous substitutions are not relevant for these two amino acids.

The matrix **R**^(*N*)^ of (15) can be expressed as the following block matrix.

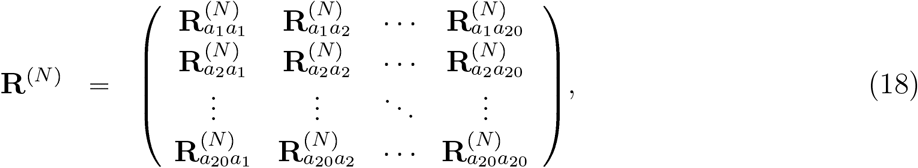

where the submatrix 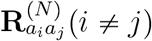 is composed of entries *s*_*a*(*r*)*a*(*s*)_*π*_*s*_ such that *a*(*r*)= *a*_*i*_ and *a*(*s*) =*a*_*j*_.

That is,

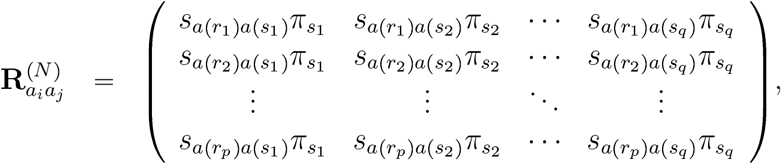

where amino acid *a*_*i*_ is encoded by *p* different codons and amino acid *a*_*j*_ is encoded by *q* different codons so that *a*(*r*_1_)= *a*(*r*_2_) =· · ·= *a*(*r*_*p*_)= *a*_*i*_ and *a*(*s*_1_) =*a*(*s*_2_) =· · · =*a*(*s*_*q*_) =*a*_*j*_. The block matrix 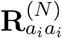 in the diagonal position of (18) is the following diagonal matrix with all off-diagonal entries being zero,

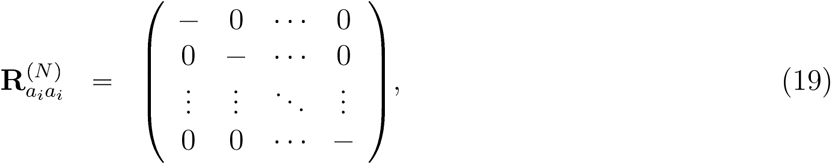

with diagonal elements being the negative sum of all entries in the same row of 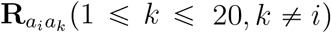.

### Transition probability matrices of new model

The transition probability matrix for time *t* that emerges from the rate matrix of (15) can be obtained with matrix exponentiation, exp{*t***R**^(*NS*)^}. We found that multiplication of **R**^(*N*)^ and **R**^(*S*)^ is commutative (see APPENDIX),

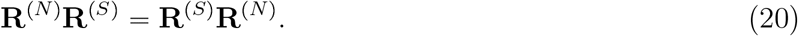

This commutative property makes it possible to factor the transition probability matrix into non-synonymous and synonymous parts (e.g., Golub and Van Loan 1996),

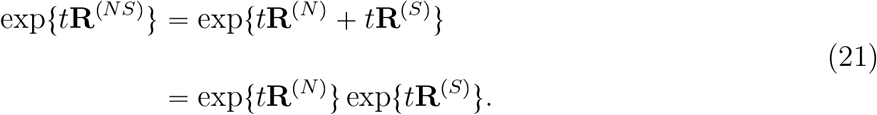

The transition probabilities of exp{*t***R**^(*N*)^} can be obtained by assigning *ρ =* 0 in (6),

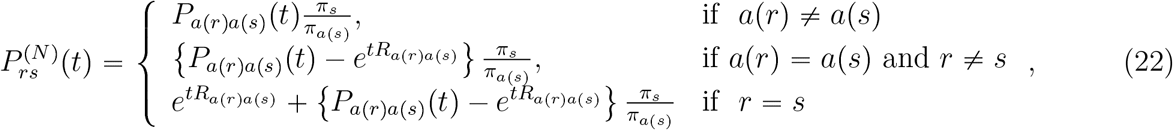

where *P*_*a*(*r*)*a*(*s*)_(*t*) and *R*_*a*(*r*)*a*(*s*)_(*t*) are derived from the 20-state amino acid model of (3). Because we use ECs that are normalized as in (4) and we do not normalize **R**^(*NS*)^, the parameter *t* is interpreted as ‘the expected number of nonsynonymous (amino acid) substitutions per codon site’.

The transition probabilities of exp{*t***R**^(*S*)^} can be obtained by exponentiating the diagonal block matrices of (17),

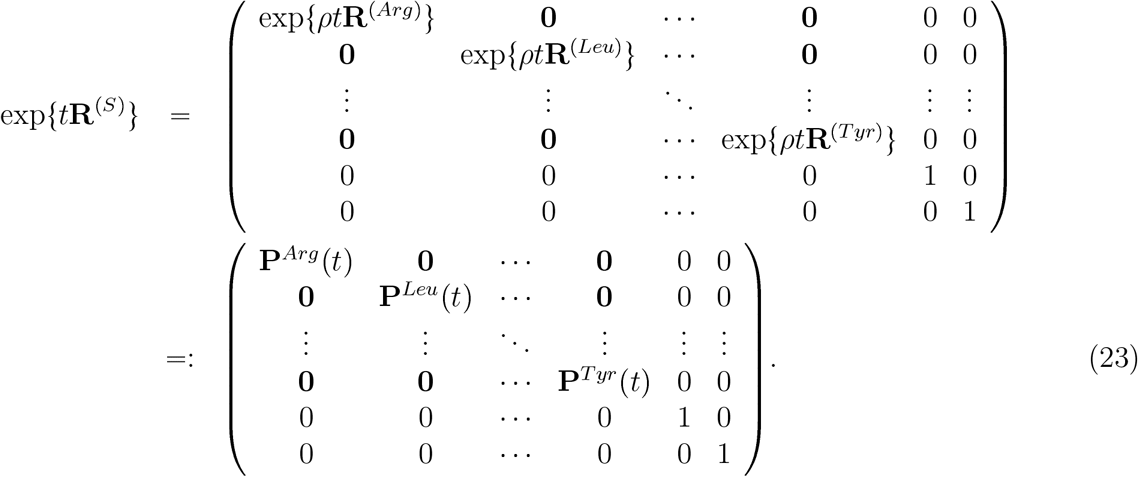

For the standard codon table, there are nine {2 ×2} diagonal block matrices in (23) and these have an analytic form for their transition probabilities. However, the matrices with a dimension that is greater than two generally do not have a closed analytic form. For those matrices, transition matrices can be obtained numerically. Denote 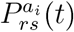 as the transition probability from codon *r* to codon *s* during time *t* within the same amino acid category *a*_*i*_. That is, 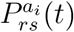 is the (*r, s*) entry of the block matrix 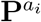 (*t*) in (23). Using the transition probabilities from exp{*t***R**^(*S*)^}, the transition probabilities of exp{*t***R**^(*NS*)^} are obtained via matrix multiplication exp{*t***R**^(*N*)^} exp{*t***R**^(*S*)^} as

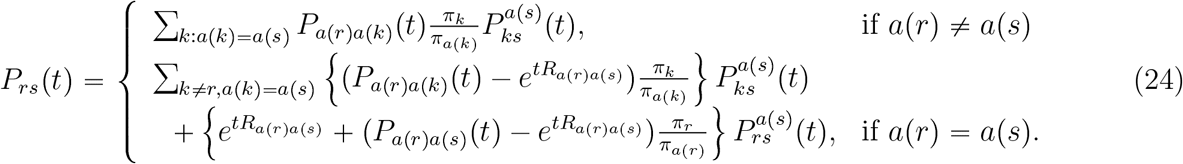

Time reversibility within each synonymous block matrix 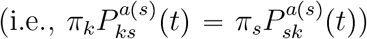 allows (24) to be simplified to

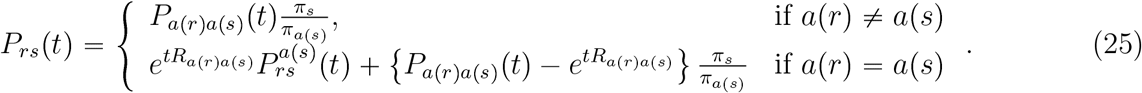

For the *a*(*r*) = *a*(*s*) case in (25), the 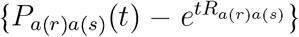 term is the probability that there has been at least one nonsynonymous change but the ending state codes for amino acid *a*(*s*). When this happens, the probability of ending with codon *s* given the ending amino acid *a*(*s*) is *π*_*s*_/*π*_*a*(*s*)_. The 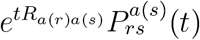 term for the *a*(*r*)= *a*(*s*) case in (25) is the probability of no nonsynonymous changes multiplied by the probability of ending with codon *s* when there have been no nonsynonymous changes. We note that this interpretation of the transition probabilities in (25) is reminiscent of the transition probability interpretation of the Felsenstein 1984 model (PHYLIP computer package, versions 2.6 and later; Felsenstein 1989) that is given in Thorne et al. (1992).

In fact, the SK-P1 transition probabilities of (6) are a special case of (25). For the SK-P1 model, the **R**^(*S*)^ matrix has transition probabilities with a similar form to those of the F81 DNA model (Felsenstein 1981),

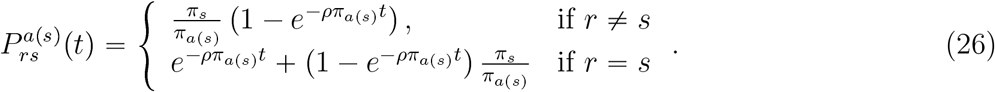

Replacing 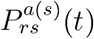 of (25) with (26) leads to (6).

Relative to conventional codon substitution models, the transition probabilities of (25) have computational advantages. Although *ρ* is a parameter of the rate matrix in (11), diagonalization of the rate matrix is not required for updating *ρ* during parameter optimization due to the special structure of synonymous blocks of (17). Furthermore, when parameters within *λ*_*S*_(*r, s*) are updated during optimization, diagonalization of the rate matrix can be done in the smaller diagonal blocks of (17). Although computational efficiency depends on various factors such as programming language, compiler, computer architecture, etc., the structure of (17) may have potential advantages.

Seo and Kishino (2008) noted that, when *ρ=* ∞, the SK-P1 model is reduced to the SK-P0 model (i.e., the product of the SK-P0 factor of (8) and the likelihood of a 20-state amino acid model). The same model reduction holds when *ρ=* ∞ for the combination of *λ*_*N*_ (*r, s*) ≡1 and *λ*_*S*_(*r, s*) ≡GTR ⊕ *ϵ* in (11).

### Equivalence classes for synonymous substitution

For the model that combines *λ*_*N*_ (*r, s*) ≡ 1 and *λ*_*S*_(*r, s*) ≡ GTR in (11), the synonymous substitution rates among arginine codons that are shown in (16) form an irreducible Markov chain. We will use ‘↔’ to indicate that two codons eventually communicate with each other via exclusively synonymous changes (e.g., TCT ↔ TCC) and we will use ↮ otherwise. The synonymous substitution rates among serine codons for the *λ*_*N*_ (*r, s*) ≡1 and *λ*_*S*_(*r, s*) ≡GTR model are

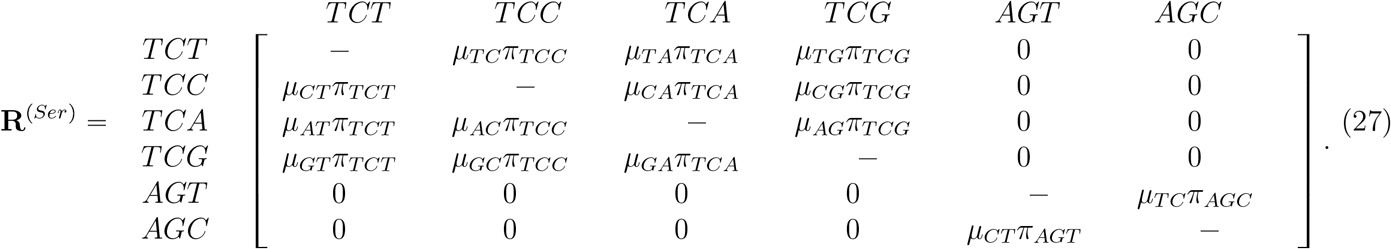

For this rate matrix, the synonymous Markov process within the serine group is not irreducible because codons in the {TCT, TCC, TCA, TCG} group cannot be transformed to codons in the {AGT, AGC} group via exclusively single position synonymous changes (e.g., Averof et al. 2000; Rogozin et al. 2016). Similarly, codons in the latter group cannot be transformed into codons in the former group via exclusively synonymous changes.

For the convenience of further discussion of the synonymous serine block of (27), we assign integers 1–6 to the corresponding six codons. Then, codons 1-4 comprise one equivalence class and codons 5–6 comprise the other. When *ρ* → ∞ and *λ*_*S*_(*r, s*) ≡GTR, the transition probabilities within the serine group approach

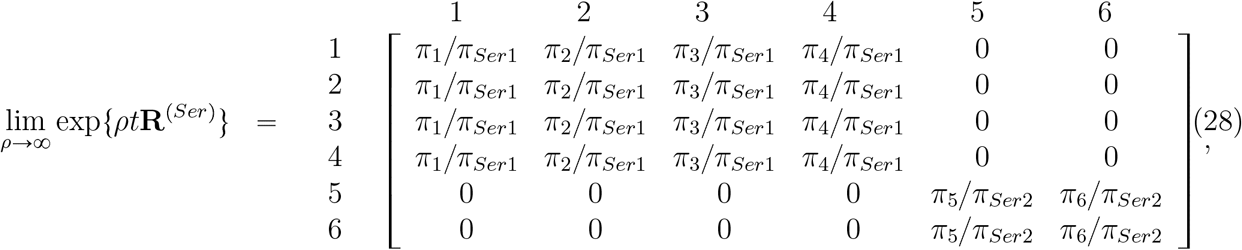

where *π*_*Ser*1_ and *π*_*Ser*2_ represent the sum of codon frequencies for each equivalence class. That is, 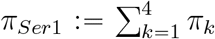 and 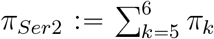. In this model setting with *λ*_*N*_ (*r, s*) ≡1, *λ*_*S*_(*r, s*) ≡GTR and *ρ=* ∞, the transition probabilities are

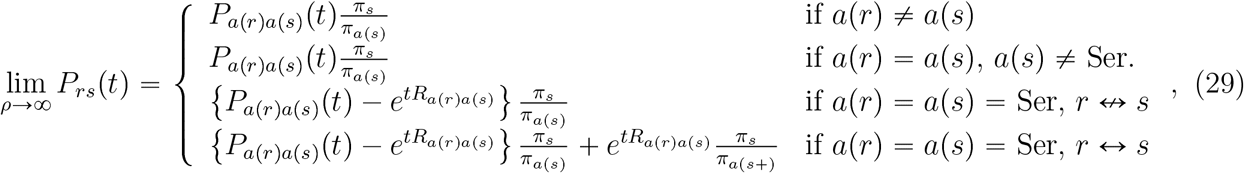

where *π*_*a*(*s*+)_ is the sum of codon frequencies within the same equivalence class as codon *s* (e.g.,*π*_*Ser*1_ or *π*_*Ser*2_ in (28)).

Equation (29) shows that the transition probabilities do not reduce to the transition probabilities of (7) and therefore (8) does not hold. That is, the transition probability is not factored into a simple form of amino acid transition probability and relative codon ratio, and therefore the SK-P0-like factor is not represented in a simple form. In contrast, all Ser codons communicate with each other when *ϵ* > 0 with *λ*_*N*_ (*r, s*) ≡ 1 and *λ*_*S*_(*r, s*) ≡ GTR⊕ *ϵ* in (11). This means that (7) and (8) hold for this ‘⊕*ϵ*’ model when *ρ=* ∞. Lumpability (Kemeny and Snell 1976) has been investigated in phylogenetic analysis (Vera-Ruiz et al. 2014, 2022). The models of (11) with *λ*_*N*_ (*r, s*) ≡ 1 satisfy the necessary and sufficient conditions for lumpability: lumping 61-state codons to 20-state amino acids. That is, the sum of transition probabilities from a starting codon to ending codons encoding the same amino acid pair is identical for all starting codons encoding the same amino acid. We note that (29) satisfies lumpability conditions, but that (29) does not lead to (7) and (8). Therefore, (29) does not lead to simple factored form of transition probabilities and likelihood. For the relationship of (7) and (8) to hold, we need an additional condition beyond lumpability: irreducibility of the Markov process of synonymous substitution.

### Eigenvalues and eigenvectors of new rate matrix

Although transition probabilities of (25) can be obtained from (20 – 23), they also can be obtained with direct exponentiation of **R**^(*NS*)^. To exponentiate **R**^(*NS*)^, we can diagonalize **R**^(*NS*)^ by obtaining matrices **U, D** and **V** such that

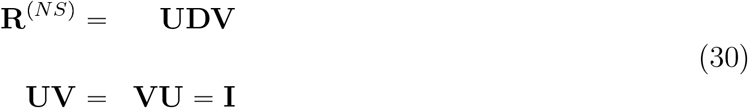

where **D** is a diagonal matrix whose entries are eigenvalues of **R**^(*NS*)^ and the column vectors of **U** are the corresponding eigenvectors. When **R**^(*NS*)^ is separated into the sum of **R**^(*N*)^ and **R**^(*S*)^, we can represent the eigenvalues and eigenvectors of **R**^(*NS*)^ with those of **R**^(*N*)^ and **R**^(*S*)^, as is described in APPENDIX. The theory is a generalization of Seo and Kishino (2008) in which **R**^(*S*)^ has identical ECs for all codon pairs as shown in (5). We show how to derive **U, D** and **V** when **R**^(*S*)^ takes a more complicated form.

### Generality of recoding approach

Our focus is on recoding codons into amino acids, but the approach described here can be employed for other situations where characters in a larger state space can be mapped into a smaller state space. The transition probabilities of (25) apply when *r* and *s* represent states from the larger state space and when *a*(*r*) and *a*(*s*) represent states from the smaller (grouped) state space. The key point is that *λ*_*N*_ (*r, s*) ≡1 for all *r* and *s* induces lumpability so that transition probabilities on the larger state space can be calculated by distinguishing between within-group and between-group transitions. Here, *P*_*a*(*r*)*a*(*s*)_ (*t*) and 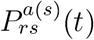represent between-group and within-group transition probabilities respectively.

### RY recoding

In the 4-state HKY (Hasegawa et al. 1985) and TN93 (Tamura and Nei 1993) DNA substitution models, transversion substitutions are analogous to nonsynonymous (i.e., between-group) substitutions and transition substitutions are analogous to synonymous (i.e., within-group) substitutions. Although the framework is also applicable to the TN93 model, we exemplify it via the HKY model. The HKY rate matrix can be represented as

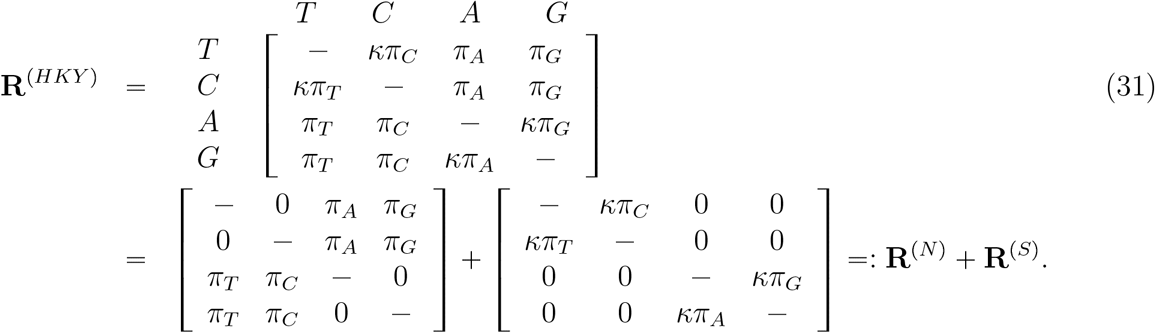

With this representation, the HKY transition probabilities of (25) have

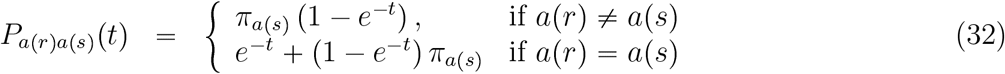

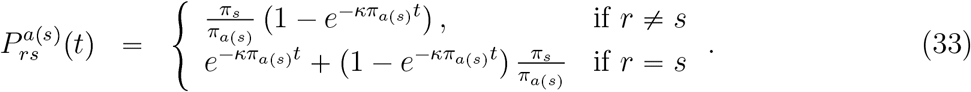

Employing this perspective, conventional RY-coding implicitly assumes that transitions are com-pletely saturated (i.e., *κ* =∞). In this case, the SK-P0-like factor has a similar form to the SK-P0 factor of (8) and is the product of *π*_*r*_/*π*_*a*(*r*)_ where *π*_*r*_ is the stationary frequency of nucleotide *r* and *π*_*a*(*r*)_ is the stationary frequency of either purines or pyrimidines depending on whether *r* is a purine or pyrimidine. This SK-P0-like factor should be multiplied by the RY-coding likelihood when comparing recoded 2-state models with 4-state models.

### Recoding individual amino acids into groups of amino acids

The transition probabilities of (25) also apply to recoding individual amino acids into groups. In this situation, *r* and *s* from (25) would represent amino acids whereas *a*(*r*) and *a*(*s*) would represent the recoded (grouped) states. Consider the 6 groups of amino acids that were proposed by Dayhoff et al. (1978): *{*A,G,P,S,T*}, {*D,E,N,Q*}, {*H,K,R*}, {*F,Y,W*}, {*I,L,M,V*}*, and *{*C*}*. A {6 × 6} between-group rate matrix could be parameterized by

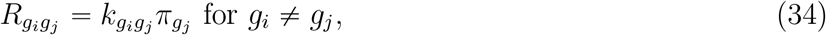

where 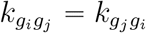 is an EC between groups *g*_*i*_ and *g*_*j*_ and 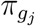 is the stationary frequency of group *g*_*j*_. With this {6 × 6} rate matrix, in a similar way to (11), a {20 × 20} rate matrix can be defined as

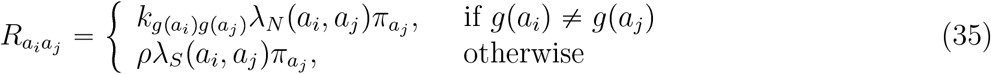

where *g*(*a*_*i*_) represents the group to which amino acid *a*_*i*_ belongs. Here, the *λ*_*i*_(*a*_*i*_, *a*_*j*_) functions define substitution within and between groups in a similar way to (12). In (34) and (35), we assume 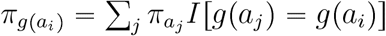 and *λ*_*S*_(*a*_*i*_, *a*_*j*_) =*λ*_*S*_(*a*_*j*_, *a*_*i*_) for all *a*_*i*_ ≠*a*_*j*_ with *g*(*a*_*i*_) =*g*(*a*_*j*_). Whereas nucleotide substitution patterns are considered in (12), physical and/or chemical properties can be considered in defining *λ*_*S*_(*a*_*i*_, *a*_*j*_) functions. Then, when *λ*_*N*_ (*a*_*i*_, *a*_*j*_) ≡1, (25) represents transition probabilities among 20 amino acids that would depend on the between-group and within-group transition probabilities. There are multiple ways to specify 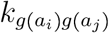and *λ*_*S*_(*a*_*i*_, *a*_*j*_). For example, 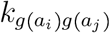 can simply have 14 (= (36− 6)/2− 1) free parameters (e.g., Feuda et al. 2017). Alternatively, the 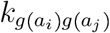 terms could depend on exchange coefficients from an empirically-derived amino acid replacement model (e.g., Kosiol et al. 2004; Susko and Roger 2007),

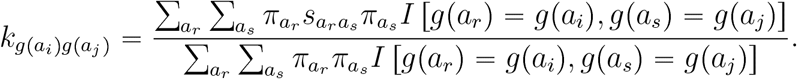

In this alternative case, there are no free parameters because the 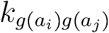 values would result from the predetermined 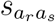values of the amino acid model.

Assuming the *λ*_*S*_(*a*_*i*_, *a*_*j*_) specifications yield irreducible within-group Markov processes, the *ρ* =∞ setting would allow a 20-state and a 6-state model to be compared by multiplying the 6-state likelihood by the SK-P0-like factor

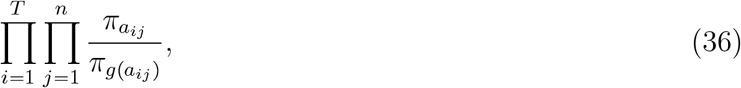

where 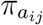is the stationary frequency of the amino acid at site *j* of the *i*th taxon and 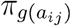 is the frequency of the group to which *a*_*ij*_ belongs. Furthermore, if codon states (*c*_*ij*_’s) are known, the following SK-P0-like factor

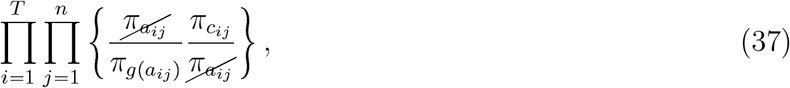

facilitates comparison of 6-state models and 61-state codon models. Alternatively, if codon states (*r* and *s*) are known, (35) can be modified as

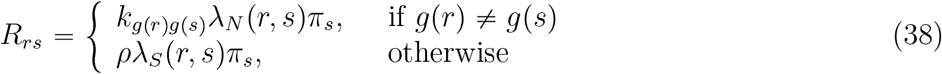

in order to improve the performance of a 6-state amino acid model in the space of 61-state codons. In addition to the 6 amino acid groups suggested by Dayhoff et al. (1978), there are other reasonable ways of grouping amino acid types (e.g., Susko and Roger 2007; Kosiol et al. 2004). We note that the likelihood scores from different grouping schemes are not directly comparable even when the total number of groups is the same in different grouping schemes. However, the approach here allows for different grouping schemes to each be converted to 20-state amino acid models or 61-state codon models via SK-P0-like multiplying factors. The likelihoods that result in the recoded 20-state or 61-state character space can be compared with conventional likelihood-based measures.

## RESULTS and DISCUSSION

### Model comparison with empirical data

We report on analyses of protein-coding genes from (1) 69 mammalian mitochondrial genomes (Nikaido et al. 2003); (2) 33 human mitochondrial genomes (Ingman et al. 2000; Seo and Kishino 2009); and (3) ATP8 genes of 69 mammals. Data Set 1 consists of sequences with moderate amounts of nonsynonymous change and large amounts of synonymous change. Data Set 2 represents 33 closely related mitochondrial genomes that were selected by Seo and Kishino (2009) from the original Ingman et al. (2000) data set. Very little nonsynonymous or synonymous change differentiates the sequences of Data Set 2. Data Set 1 and Data Set 2 are both comprised of multiple protein-coding sequences that have been concatenated for our analyses. Data Set 3 is a subset of Data Set 1. Whereas Data Set 1 has sequences from 12 protein-coding genes and a concatenated length of 3705 codons, Data Set 3 represents only the ATP8 gene that has a length of 70 codons. We adopted the codon table for mammalian mitochondria in which there are 60 sense codons. Our theory equally applies to a codon table whose character dimension is different from 61.

All analyses with codon substitution models were done with prespecified tree topologies. For Data Set 1, we estimated the maximum likelihood (ML) tree topology with the IQ-TREE program (Nguyen et al. 2015) by adopting the mtREV amino acid replacement model (Adachi and Hasegawa 1996). We used empirically-derived amino acid frequencies (Cao et al. 1994), and discretized-gamma rate heterogeneity among sites (RHAS; Yang 1994) with 5 rate categories (mtREV+F+G5). For Data Set 2, we adopted the tree topology that was inferred and used in previous studies (Ingman et al. 2000; Seo and Kishino 2009). For Data Set 3, we used the tree topology that was inferred with Data Set 1.

All codon substitution analyses allowed RHAS via a 5-category discretized gamma distribution (Yang 1994). The relative rate for a codon location in one of the 5 categories was applied to both synonymous and nonsynonymous changes. For stationary frequencies of codons and amino acids, we used empirical frequencies obtained from the data being analyzed.

The three data sets were each analyzed with four previously proposed models (GY94, SK-P0, SK-P1, and SK-P2) and with seven parameterizations that are cases of the codon substitution rates of (11). Table 1 details all 11 codon substitution models that were explored.

Let us consider the SK-P1 parameterization and the new parameterizations that will be referred to as P2 and P6 as follows:

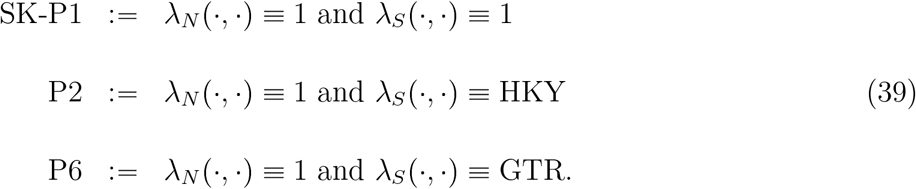

The P2 name stems from the fact that it has 2 parameters (*ρ* and *κ* from *λ*_*S*_(·, ·)) that are estimated by maximum likelihood in the rates of (11). Similarly, P6 has 6 parameters: *ρ* and five *µ*_*ij*_ parameters of (13). For identifiability, we set one of the *µ*_*ij*_ parameters to one.

The P2 and P6 implementations used 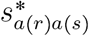 values that were synchronized with amino acid replacement rates as in (14). Because *λ*_*N*_ (·, ·) 1 for P2 and P6, this synchronization amounted to setting 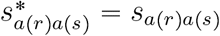.

For comparison of P6 with P2, an LRT is applicable. The critical region of the log-likelihood difference with a 5 % significance level is (4.744, ∞). For all three data sets, the log-likelihood score of P6 is greater than that of P2. However, their difference is located within the critical region for Data Sets 1 and 2, whereas it is not for Data Set 3. Due to short sequences, Data Set 3 does not seem to have accumulated enough synonymous substitution to distinguish the GTR from the HKY model. Both P2 and P6 models show greater log-likelihood scores than SK-P1 in all three Data Sets. Due to the structure of the ser block in (27), an LRT is not applicable for this comparison, but AIC can be used instead for model comparison. By involving appropriate 4-state nucleotide substitution models, we could greatly improve the model fit of P2 and P6. Although incorporation of the GTR model into a codon model may improve model fit, there is a resulting computational cost associated with having to optimize additional parameters.

We note that modifying the *λ*_*N*_ (·, ·) 1 setting may substantially improve model-fit. Let us consider the SK-P2 parameterization and three more parameterizations with/without synchronization of (14), which will be referred to as P3a, P3b and P3c as follows:

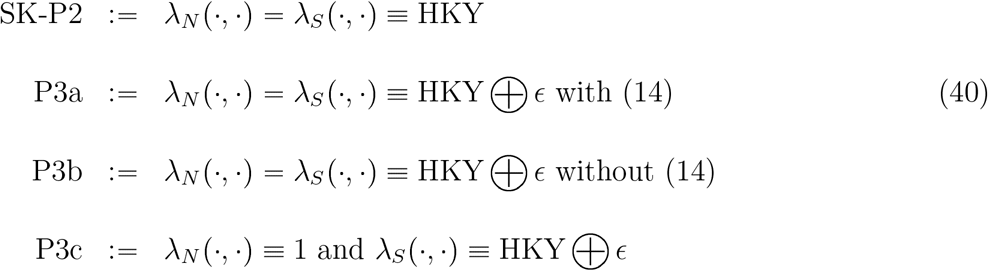

where the *κ* parameter of the HKY model is shared in both *λ*_*N*_ (·, ·) and *λ*_*S*_(·, ·). In the P3a and P3b models, *ϵ* of (12) is also shared in both functions. P3c, which assumes *λ*_*N*_ (·, ·) 1, nests P2 of (39) by adding *ϵ* to synonymous substitution. Whereas synchronization of 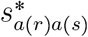 values of (14) is adopted in P3a, it is not in P3b.

Although P3c nests P2, the LRT statistic asymptotically follows a 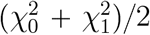 distribution under the null hypothesis (Self and Liang 1987) because the value *ϵ* = 0 for P2 is on the boundary of the parameter space. Therefore, the critical region of the log-likelihood difference with a 5 % significance level is (1.353, ∞). Whereas the log-likelihood difference of Data Set 1 is within this critical region, those of the other two data sets are not. This may imply that the effect of multiple nucleotide changes is minor in the codon substitution modeling when sequences are not divergent and/or sequences are relatively short. Comparison among P3a, P3b and P3c can be done with AIC. For the three data sets, either P3a is better than P3b (Data Sets 1 and 3) or P3b is better than P3a (Data Set 2). Whereas both P3a and P3b are better than P3c in Data Sets 2 and 3, P3c is better than P3b in Data Set 1, which implies that relaxing the *λ*_*N*_ (·, ·) 1 assumption may not necessarily increase model-fit if it is combined with synchronization of (14).

We considered two parameterizations, which will be referred to as P5a and P5b, by relaxing the assumption of sharing *κ* and *ϵ* parameters in P3a and P3b as follows:

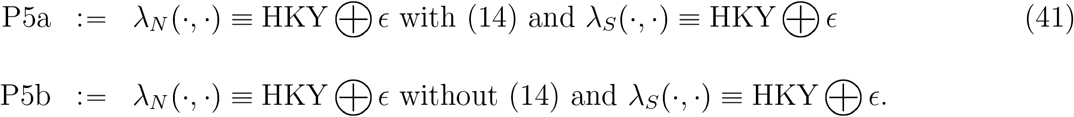

Here, synonymous and nonsynonymous substitutions have separate *κ* and *ϵ* parameters.

Comparison of P5a and P5b with P3a and P3b can be done with an LRT, and the critical region of the log-likelihood difference with a 5 % significance level is (2.996, ∞). Except for the case of P3b vs. P5b with Data Set 2, the log-likelihood differences are significant. This implies that allowing different substitution patterns for synonymous and nonsynonymous change is more realistic.

For the comparatively divergent Data Set 1, we found P5a is the best among the 11 evaluated models on the basis of AIC scores. The amino acid model, which is denoted as SK-P0, is worse than the other 61-state codon models. P3c is significantly better than P2. The estimated synonymous *ϵ* under the P3c model for Data Set 1 was 0.00439. Although this value is only slightly above 0, it causes a substantial log-likelihood increase compared to the P2 model. This implies that allowing multiple nucleotide changes is a more realistic model assumption for divergent sequences.

For the comparatively less divergent Data Set 2, the SK-P2 model is the best. SK-P2 being better than GY94 suggests that the incorporation of 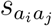 values in (9) is worthwhile. Due to the limited amount of divergence, the nonsynonymous substitution pattern does not seem to be congruent with the pattern of the provided amino acid model, which explains why P3a and P5a synchronizing 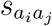 values are worse than unsynchronizing P3b and P5b. P5a being worse than SK-P2 may imply that synchronization of (14) worsens model fit when conserved protein sequences are not consistent with the change pattern of the given amino acid model. Based on AIC scores, the P2 model is better than P3c, which may imply that assuming *ϵ* 0 in (12) is more realistic for Data Set 2.

As is the case for Data Set 1, the P5a model is the best for Data Set 3. This suggests that empirical information from an amino acid model can be useful for divergent sequences in spite of their short length. However, it is notable that SK-P2 is worse than GY94 for Data Set 3. Presumably, sequences that are too short may not be informative enough to compare substitution models properly. Alternatively, even if 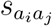 values are incorporated in SK-P2, a subtle difference in how they are incorporated may affect relative superiority/inferiority of competing models especially when sequences are very short as in Data Set 3.

### Tree topology comparison with different models

We note that model selection affects not only the optimal tree topology but also relative superiority or inferiority among competing tree topologies. As an example, we consider the analysis of 69 mammalian protein-coding sequences (Data Set 1 in Table 2).

**Table 2:** Log-likelihood scores under 11 models. #1 mammalian MT, #2 Human MT, #3 mammal ATP8.

| Data | Total | Nuc. |  |  |  |  |
| set | taxa | length | Model | LnL | # param | AIC |
|---|---|---|---|---|---|---|
| 1 | 69 | 11115 | GY94 | -347261.80 | 3+135 | 694799.60 |
|  |  |  | SK-P0 | -366035.36 | 1+135 | 732342.72 |
|  |  |  | SK-P1 | -348345.58 | 2+135 | 696965.16 |
|  |  |  | SK-P2 | -342337.95 | 3+135 | 684951.90 |
|  |  |  | P2 | -339666.77 | 3+135 | 679609.54 |
|  |  |  | P3a | -337610.52 | 4+135 | 675499.02 |
|  |  |  | P3b | -341119.29 | 4+135 | 682517.58 |
|  |  |  | P3c | -339648.12 | 4+135 | 679574.24 |
|  |  |  | P5a | -337594.82 | 6+135 | 675471.64 |
|  |  |  | P5b | -338170.90 | 6+135 | 676623.80 |
|  |  |  | P6 | -339525.69 | 7+135 | 679335.38 |
| 2 | 33 | 10845 | GY94 | -15942.89 | 3+63 | 32017.78 |
|  |  |  | SK-P0 | -122292.09 | 1+63 | 244712.18 |
|  |  |  | SK-P1 | -16227.93 | 2+63 | 32585.86 |
|  |  |  | SK-P2 | -15915.38 | 3+63 | 31962.76 |
|  |  |  | P2 | -16052.84 | 3+63 | 32237.68 |
|  |  |  | P3a | -15932.25 | 4+63 | 31998.50 |
|  |  |  | P3b | -15914.97 | 4+63 | 31963.94 |
| Data | Total | Nuc. |  |  |  |  |
| set | taxa | length | Model | LnL | # param | AIC |
|  |  |  | P3c | -16052.69 | 4+63 | 32239.38 |
|  |  |  | P5a | -15922.82 | 6+63 | 31983.64 |
|  |  |  | P5b | -15913.94 | 6+63 | 31965.88 |
|  |  |  | P6 | -16007.83 | 7+63 | 32153.66 |
| 3 | 69 | 210 | GY94 | -8327.94 | 3+135 | 16931.88 |
|  |  |  | SK-P0 | -9113.68 | 1+135 | 18499.36 |
|  |  |  | SK-P1 | -8429.65 | 2+135 | 17133.30 |
|  |  |  | SK-P2 | -8364.34 | 3+135 | 17004.68 |
|  |  |  | P2 | -8330.76 | 3+135 | 16937.52 |
|  |  |  | P3a | -8212.57 | 4+135 | 16703.14 |
|  |  |  | P3b | -8248.84 | 4+135 | 16775.68 |
|  |  |  | P3c | -8330.46 | 4+135 | 16938.92 |
|  |  |  | P5a | -8201.77 | 6+135 | 16685.54 |
|  |  |  | P5b | -8238.69 | 6+135 | 16759.38 |
|  |  |  | P6 | -8327.67 | 7+135 | 16937.34 |

For each of the 10 tree topologies (T1 – T10) that Nikaido et al. (2003) considered and also for our ML tree (T11), we calculated maximum log-likelihood scores. Then, using sitewise log-likelihood scores, we performed the Shimodaira-Hasegawa (SH) test (Shimodaira and Hasegawa 1999) under the mtREV amino acid model, P5a, and GY94 codon models (Table 3).

**Table 3:** Shimodaira-Hasegawa test for 11 trees.

| Tree # | mtREV |  | P5a |  | GY94 |  |
|---|---|---|---|---|---|---|
|  | Δloglike | p-value | Δloglike | p-value | Δloglike | p-value |
| T1 | 20.1 | 0.528 | 6.6 | 0.799 | - | 0.965 |
| T2 | 48.9 | 0.155 | 39.7 | 0.305 | 42.0 | 0.332 |
| T3 | 25.8 | 0.448 | 15.9 | 0.630 | 13.4 | 0.707 |
| T4 | 68.0 | 0.063 | 74.0 | 0.074 | 85.4 | 0.066 |
| T5 | 42.4 | 0.251 | 56.4 | 0.167 | 75.6 | 0.085 |
| T6 | 24.7 | 0.445 | 47.6 | 0.234 | 90.2 | 0.048 |
| T7 | 43.9 | 0.229 | 71.7 | 0.101 | 115.2 | 0.019 |
| T8 | 41.4 | 0.251 | 62.6 | 0.150 | 105.9 | 0.028 |
| T9 | 26.1 | 0.440 | 43.4 | 0.278 | 90.2 | 0.057 |
| T10 | 30.6 | 0.373 | 51.6 | 0.209 | 101.0 | 0.034 |
| T11 | - | 0.983 | - | 0.933 | 6.1 | 0.840 |

The best tree topology is T11 under both mtREV and P5a models. However, inferiority of the other 10 competing tree topologies is not significant although the test statistic of T4 is near the boundary (Table 3). In contrast, the best tree topology is T1 under the GY94 model and p-values of T6–T8 and T10 are less than 5% although they are not highly significant, which implies that we may make a different conclusion under the GY94 model and under mtREV and P5a models. We note that the results of topology comparison can be variable and a good model (P5a) and a bad model (mtREV) can produce consistent results whereas an intermediate model produces a different result. Without the P5a model, it would be tempting to trust the results of the GY94 model because the GY94 model is a better fit to the data than the mtREV model.

The results of Table 3 provide us with a hint for how we should proceed with data analysis when different tree topologies are supported by recoded versus original state space models. As shown in Table 3, a conventional unrecoded model (e.g., GY94 in this case) and a recoded model (e.g., mtREV in this case) can produce different tree topologies. Furthermore, it has been observed that recoded 6-state models and 20-state amino acid models produce different tree topologies (e.g., see Foster et al. 2023). From our theory, it is always possible to reconstruct an unrecoded substitution model from the given recoded substitution model. For example, we can reconstruct unrecoded 61-state SK-P1, P2, P3 models from the given recoded 20-state amino acid models. Likewise, we can reconstruct 4-state nucleotide substitution models from the given 2-state RY-substitution models. In practice, the reconstructed model is statistically better than the given recoded model. By investigating tree topologies produced by the recoded model, the reconstructed unrecoded model and the conventional unrecoded model, and by performing a statistical comparison of these models, we will be able to more efficiently figure out the relative importance of each component of evolutionary processes during model improvement.

*Computational advantages of λ*_*N*_ (*r, s*) 1 *setting*

When measuring selection strength in different lineages or at different sites, computational time increases because we should obtain the MLE of *ω* (relative occurrence of nonsynonymous substitution) of (2) or *ρ* (relative occurrence of synonymous substitution) of (11) for each branch or site. However, computational time can be reduced when *λ*_*N*_ (*r, s*) 1 is adopted in (11).

To calculate the transition probability matrix, a first step can be to diagonalize the rate matrix (**R**) as

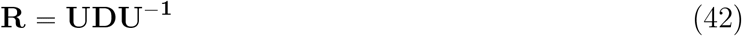

where **D** is a diagonal matrix with eigenvalues and **U** is composed of eigenvectors. Then, the transition probability matrix during time *t* is calculated as

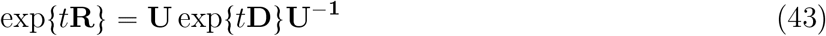

In general, when we update parameters of the rate matrix **R** and recalculate transition probabilities, we should redo the diagonalization of (42). Although *ρ* is a parameter of the {61 × 61} rate matrix of (11), we do not need to redo diagonalization of (42) for the updated *ρ* parameter. The transition probabilities among 20 amino acids, *P*_*a*(*r*)*a*(*s*)_ of (25), are free from *ρ*. Also, *ρ* is a common multiplying factor for synonymous block matrices of (17) and is free from diagonalization. In contrast to the model of (11) with *λ*_*N*_ (*r, s*) 1, the models without *λ*_*N*_ (*r, s*) 1 such as GY94 cannot have the form of (25) and diagonalization of the rate matrix is required for the updated *ω* parameter. Also, diagonalization of a {61 × 61} matrix can be numerically unstable when *ω* is very small or when entries of the rate matrix incidentally cause linear dependence among eigenvectors. The models of (11) with *λ*_*N*_ (*r, s*) 1 are relatively free from this numerical instability issue. The advantage of skipping diagonalization of the rate matrix increases when we frequently need to update *ρ* values during ML optimization in each branch or each site.

When diagonalization of the rate matrix (42) is required, such as when updating *κ* or *ϵ* parameters in block matrices of synonymous substitution (17), the models of (11) with *λ*_*N*_ (*r, s*) 1 also have potential advantages. For the case of the standard codon table, there are three {6 × 6}, five {4 × 4}, one {3 × 3}, nine {2 × 2} and two {1 × 1} synonymous block rate matrices of (17). For 11 matrices whose dimension is less than three, we can analytically calculate transition probabilities and we do not need to diagonalize these lower-rank block matrices. Instead, we only have to redo diagonalization for the other 9 block matrices whose dimension is greater than two. Although the total number of the four basic operations {+, −, ×, ÷ } for diagonalizing these 9 block matrices is substantially smaller than that of {61×61} matrix diagonalization, we should pay for the additional overhead caused by 9 function calls of diagonalization and memory access during a program run. The relative computational time for the overhead of function calls and memory access may rely on various factors, such as programming language, compiler, computer architecture, etc. Although we may not be able to expect these factors to always improve computational performance, they are potential merits of the models of (11) with *λ*_*N*_ (*r, s*) ≡ 1.

## CONCLUDING REMARKS

Here, we provided the theoretical background for a likelihood framework to compare different schemes of character recoding. Although we mainly discussed comparing a 20-state amino acid model with its derivation of 61-state codon models, our theory can be generally applied to any number of character recoding schemes. As noted by previous studies (Abadi et al. 2019) and also as supported by our analyses, the better model may produce incorrect results in some circumstances or by chance. When we mention that “model A is better than model B”, we do not imply that model A always produces a better result than model B. Instead, we imply that model A ‘on average’ has more chance to produce a better result even though it may fail occasionally. This average properly justifies our effort to find the best model among many candidates with a likelihood framework. Likelihood-based statistical analysis has a good aspect of ‘statistical consistency’ which guarantees correct results when the adopted model is correct and when the data size is infinitely large. In real data analysis, bias caused by model violation is inevitable (Box 1976). The selected better models will have more tendency toward smaller bias and therefore are expected to be closer to the property of ‘statistical consistency’.

In our model framework with *λ*_*N*_ (*r, s*) ≡ 1, we assume an identical *ρ* parameter in all diagonal block matrices of (17). We note that our theory is also valid when different *ρ*’s are assigned to different diagonal blocks. Equations (20, 25) also hold, and the eigenvalues of APPENDIX [2] are adjusted with the corresponding *ρ* parameters, while the eigenvectors remain unchanged. With different *ρ*’s, we can investigate variation in synonymous substitution rates among amino acids in future work.

Our general suggestion is to be cautious about character recoding and to quantitatively assess it. The approaches described here may be helpful in making recoding decisions. When inferences are robust to recoding, the results may be more convincing. When results differ with recoding, a quantitative comparison of the competing models might illuminate the biological underpinnings of the sources of the different results.

## ACKNOWLEDGEMENTS

The authors developed the mathematical framework and designed the matrix structures underlying Eqs. (52 – 57), following an approach analogous to that of Seo and Kishino (2008). OpenAI’s ChatGPT (GPT-6) was used as an auxiliary tool in constructing individual matrix entries consistent with the predefined structures. All AI-assisted expressions were carefully examined and verified by the authors, who take full responsibility for their mathematical validity. ChatGPT was also used for limited language editing, such as checking spelling, punctuation, grammar and article usage, while maintaining the authors’ original wording and sentence structure.

## APPENDIX

### [1] Proof of (20) : R^(*N*)^R^(*S*)^ R^(*S*)^R^(*N*)^

**R**^(*N* )^**R**^(*S*)^ is

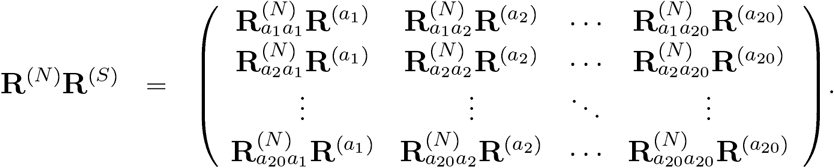

**R**^(*S*)^**R**^(*N* )^ is

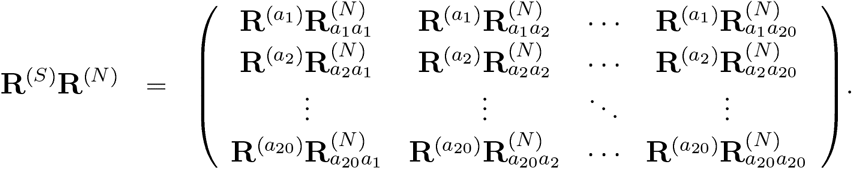

The block matrix at the *i*th row and *j*th column of **R**^(*N*)^**R**^(*S*)^ is 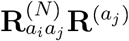. The block matrix at the *i*th row and *j*th column of **R**^(*S*)^**R**^(*N*)^ is 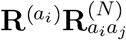 . The entries at the *r*th row and *s*th column of 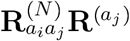 and 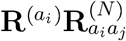 are

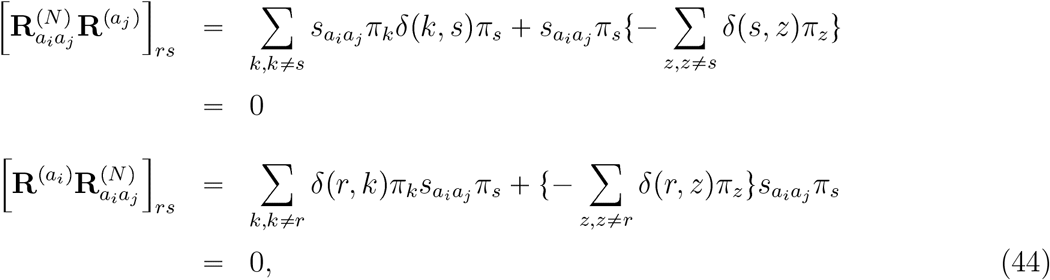

where *i* ≠*j* and symmetry of *δ*(*k, s*) is assumed in the first line: *π*_*k*_*δ*(*k, s*)*π*_*s*_ *π*_*s*_*δ*(*s, k*)*π*_*k*_. The entries at the *r*th row and *s*th column (*r* ≠*s*) of 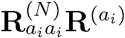 and 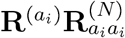 are

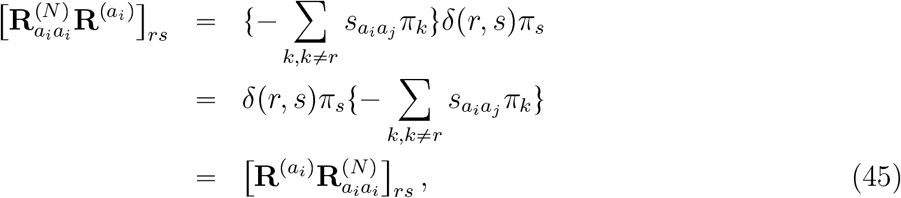

The entries at the *r*th row and *r*th column of 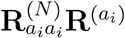 and 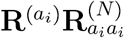 are

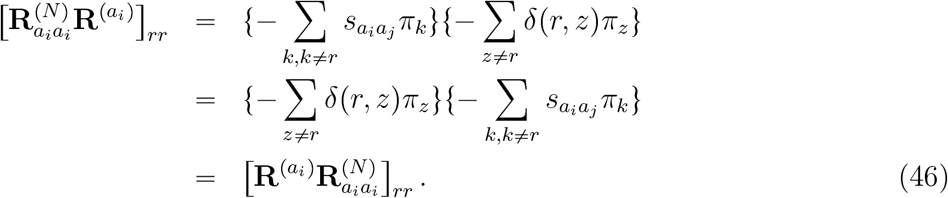

Taken together, (44) – (46) demonstrate that **R**^(*N*)^**R**^(*S*)^ = **R**^(*S*)^**R**^(*N*)^.

#### [2] Derivation of eigenvalues and eigenvectors in (30)

Because our derivation is a generalization of Seo and Kishino’s (2008), we make our mathematical notation compatible with theirs so that comparing and understanding both theories may become convenient.

For 1 ⩽ *i ⩽* 20, 1 ⩽ *j ⩽* 20, let us denote **R**_*a*_ as the 20-state amino acid rate matrix derived from (3, 4),

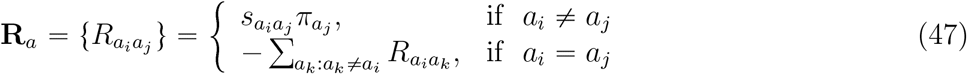

**R**_*a*_ is diagonalized as follows.

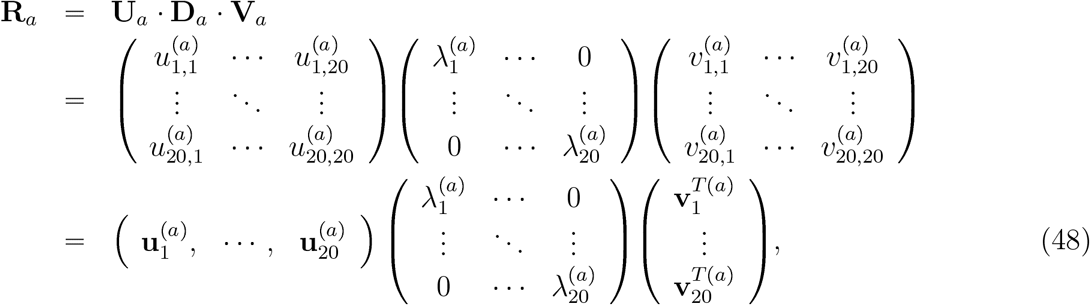

where 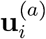 is the eigenvector of **R**_*a*_ and 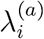 is the eigenvalue corresponding to 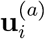. **U**_*a*_**V**_*a*_ *=* **V**_*a*_**U**_*a=*_ **I**. Due to the special structure of a continuous-time Markov process, one of the eigenvalues is zero. For convenience, we place the zero eigenvalue in the first place, i.e., 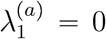. All entries of the corresponding eigenvector 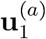 are identical. For convenience, we assume 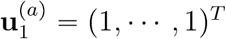.

Now, let us consider a special case of (11) where *λ*_*N*_ (*r, s*) 1 and therefore 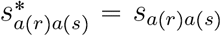. Let us denote this 61 × 61 rate matrix as **R**_*c*_. Then,

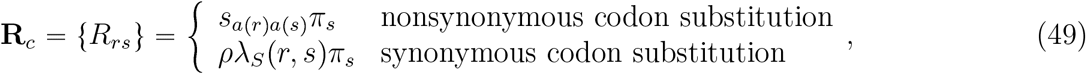

We follow Seo and Kishino (2008) in determining the order of codons within **R**_*c*_, which is briefly reviewed here. Codon ordering of **R**_*c*_ should be compatible with the amino acid ordering of **R**_*a*_. The order of amino acids can be arbitrarily determined. Once the order of amino acids is determined, the order of codons is arranged so that codons encoding the same amino acid are placed together in accordance with the amino acid order. Codon order within the same amino acid group is arbitrarily determined. For the convenience of description, we use a different amino acid order than (17). We place isoleucine (Ile) and valine (Val) and their corresponding codons (ATT, ATC, and ATA for isoleucine and GTT, GTC, GTA, and GTG for valine) in the first part of rows and columns as in Seo and Kishino (2008). Also, for convenience, we interchangeably use the integer index with the amino acid index and the codon index. For example, we interchangeably use 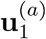 and 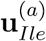.

Similar to (48), **R**_*c*_ is diagonalized as follows:

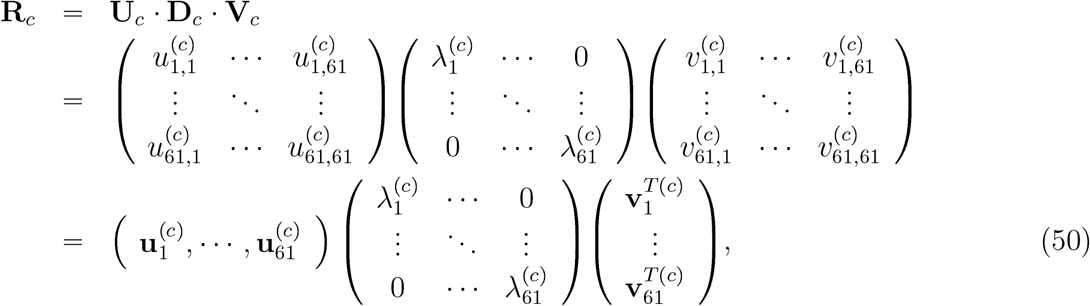

where **U**_*c*_**V**_*c*_*=* **V**_*c*_**U**_*c*_ *=* **I**.

For the SK-P1 model in which **R**_*c*_ of (49) has *λ*_*S*_(*r, s*) ≡ 1, Seo and Kishino (2008) presented eigenvalues and eigenvectors of **R**_*c*_ by using those of **R**_*a*_, *ρ*, and the frequencies of amino acids and codons. We generalize Seo and Kishino’s presentation so that it can cover the general case of *λ*_*S*_(*r, s*) ≢ 1.

Let us denote 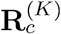 as the diagonal block matrix for amino acid *K* in (17). Then, its corresponding 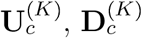 and 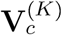 matrices should satisfy the following.

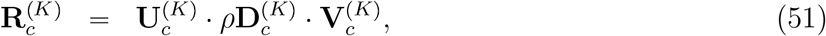

where 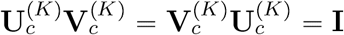. We represent the entries of 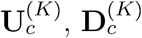 and 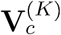 matrices with 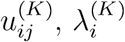 and 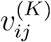. Due to the special structure of a continuous-time Markov process, one of 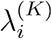 should be zero and we assume 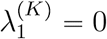 and 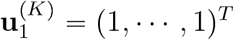.

We found that **D**_*c*_ of (50) is represented as

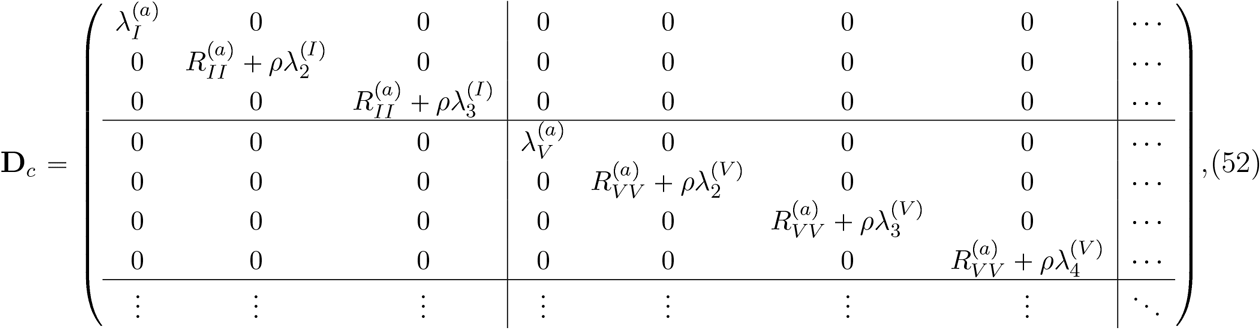

where *I* and *V* are single characters for isoleucine and valine. 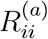 is a diagonal entry of the 20-state matrix of (47). 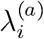 is an eigenvalue of (48) and 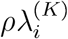 is an eigenvalue of (51).

We found that **U**_*c*_ is

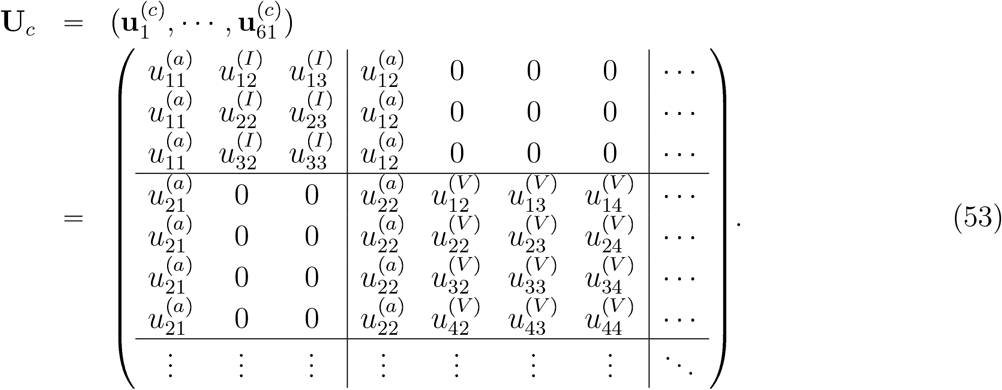

where 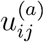 is the entry of **U** of (48) and 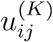 is the entry of 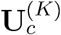 in (51).

We found that **V**_*c*_ is

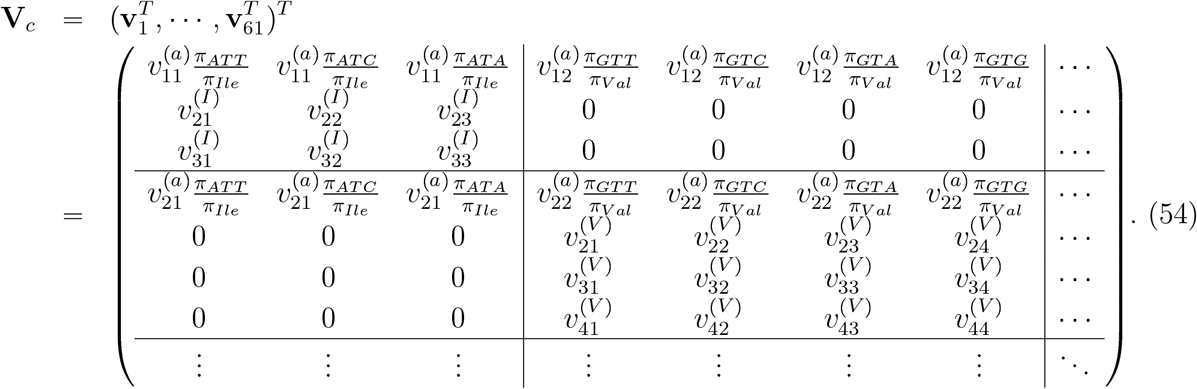

where 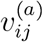 is the entry of **V** of (48) and 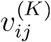 is the entry of 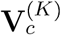 in (51).

As we showed in (27), the serine group may not be irreducible depending on the definition of *λ*_*S*_(*r, s*) in (49). In this case, we should be careful in determining the entries of **U**_*c*_, **D**_*c*_ and **V**_*c*_. To represent eigenvalues and eigenvectors of multiple equivalence classes, we replace valine with serine in the second position of the amino acid order. Then, the matrix **D**_*c*_ comprising eigenvalues is represented as

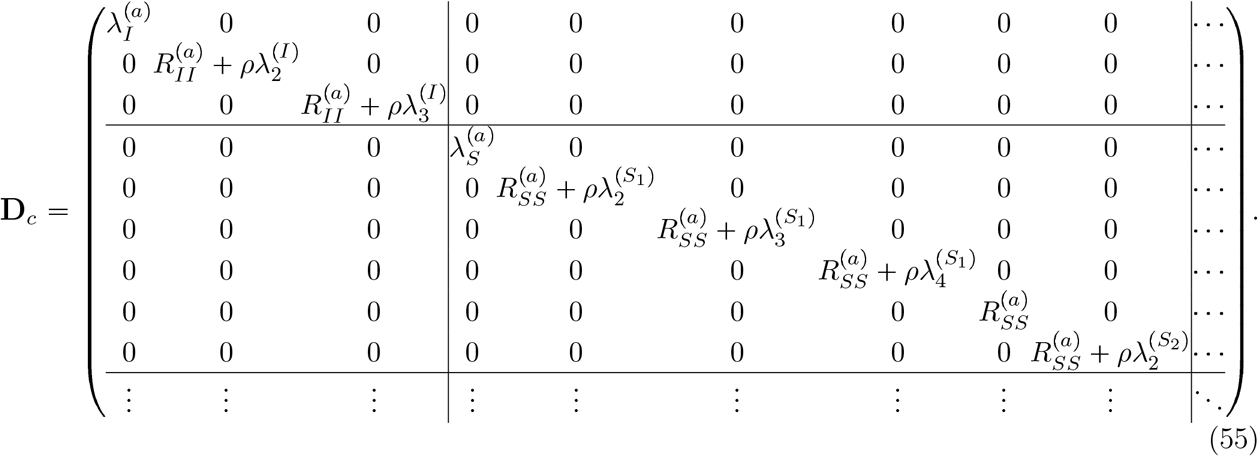

where *S*_1_ is serine encoded by the first four codons of (27) and *S*_2_ is serine encoded by the last two codons. 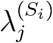 is the j*th* eigenvalue of the *S*_*i*_ equivalence class within the serine block.

The matrix **U**_*c*_ is represented as

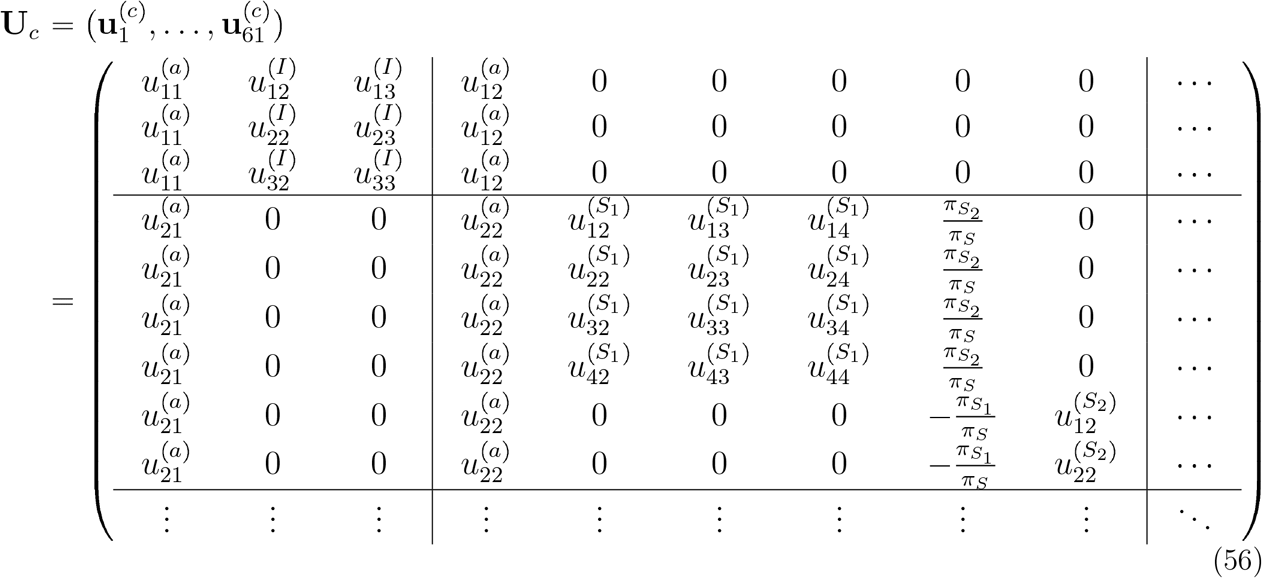

The matrix **V**_*c*_, which is the inverse of **U**_*c*_, is represented as

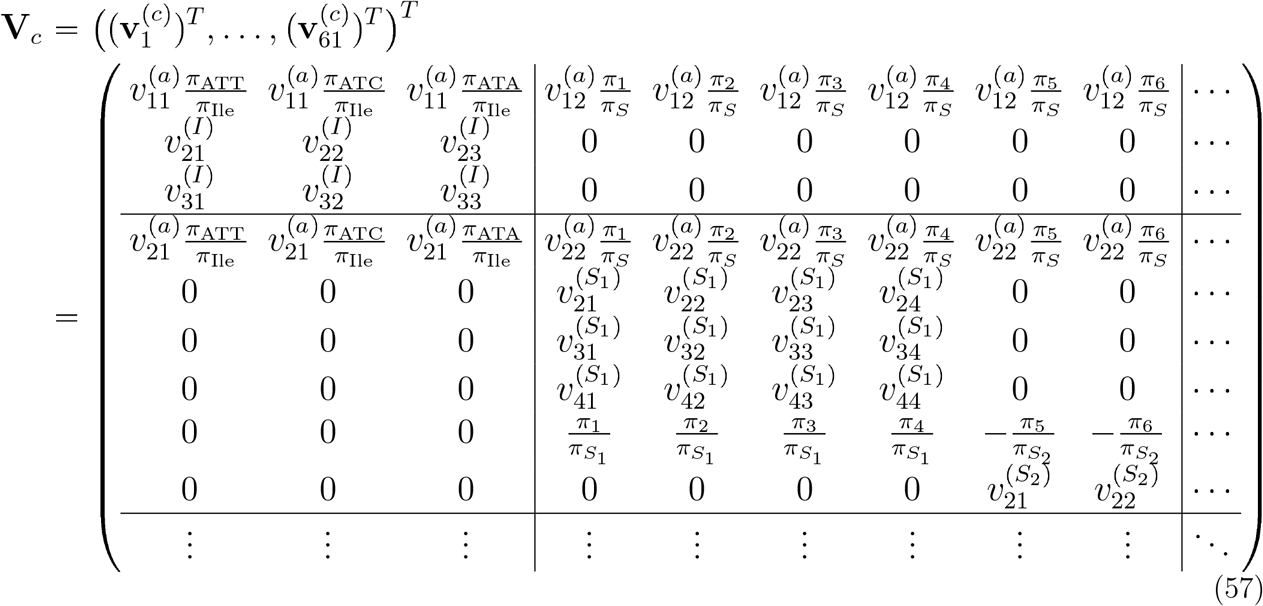

where *π*_1_, · · ·, *π*_6_ follow the notation of (28) and 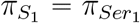 and 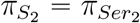.

Now, we formalize (52, 55), (53, 56) and (54, 57) respectively, by defining 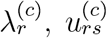 and 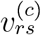 entries as follows.

For 1 ⩽*r⩽* 61, the diagonal entries of **D**_*c*_ in (55), eigenvalues 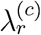, are defined as

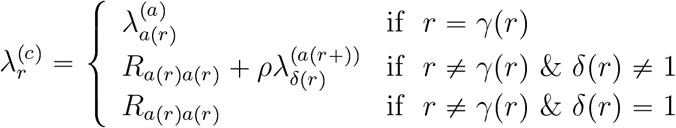

where *γ*(*r*) is the index of the first codon that encodes *a*(*r*). For example, *γ*(1) = *γ*(2) = *γ*(3) = 1 because the first codon (ATT) encoding Ile is located in the first row and column of **R**_*c*_. When Val is placed in the second amino acid position, *γ*(4) = *γ*(5) = *γ*(6) *γ*(7) = 4 because the first codon (GTT) encoding Val is located in the fourth row and column of **R**_*c*_. When serine is placed in the second amino acid position, *γ*(4) = *γ*(5) = *γ*(6) = *γ*(7) = *γ*(8) = *γ*(9) = 4. The function *δ*(*r*) indicates the integer index of *r* within the equivalence class. Because Val’s codons constitute a single equivalence class, *δ*(4) = 1, *δ*(5) = 2, *δ*(6) = 3, *δ*(7) = 4 when Val is placed in the second amino acid position. When serine is placed in the second amino acid position, *δ*(4) = 1, *δ*(5) = 2, *δ*(6) = 3, *δ*(7) = 4, *δ*(8) = 1, *δ*(9) = 2.

For 1 ⩽ *r⩽* 61, 1⩽ *s⩽* 61, the entries of the eigenvectors, 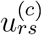, are defined as

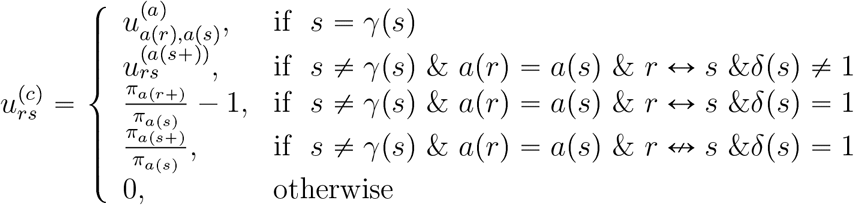

where 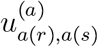 is the element of 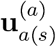 in (48). 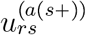 indicates the entry of the 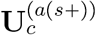 matrix within the same equivalence class as codon *s. π*_*a*(*s*+)_ is the sum of codon frequencies within the same equivalence class as *s*.

The entries of **V**_*c*_, 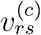, are defined as

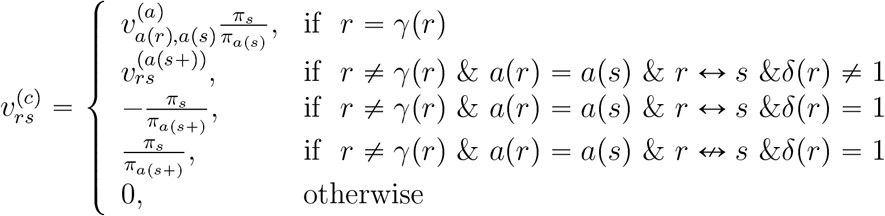

where 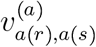 is the element of 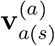 in (48). 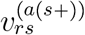 indicates the entry of the 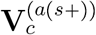 matrix within the same equivalence class as codon *s*.

The transition probabilities between codons can be calculated using the matrix exponential

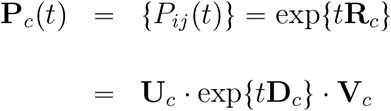

which is identical to the derivation based on (21). We note that the above-mentioned results (55 – 57) are specific for an amino acid block in which there are two equivalence classes. The pattern of (55 – 57) is different when an amino acid block contains more than two equivalence classes (data not shown). In contrast, the derivation based on (21) is applicable irrespective of the number of equivalence classes.

